# A network-based comparative method to study reptile scalation and other homologous two-dimensional patterns

**DOI:** 10.1101/2025.02.23.639519

**Authors:** Isaac W. Krone

**Affiliations:** Museum of Vertebrate Zoology, University of California, Berkeley

**Keywords:** networks, comparative methods, squamates, evolution, morphometrics, fossorial

## Abstract

From the heads of sunflowers to insect wings to the scales on the heads of reptiles, discrete, two-dimensional patterns abound in the natural world. While some of these patterns, like plant phyllotaxis and the regular scalation of snake bodies, are created by repetition following a simple algorithm, others are far more complex and idiosyncratic, containing irregular elements that nonetheless have clear homology relationships between species. Despite the importance of these patterns in species delimitation and taxonomy researchers have not developed holistic methods for comparing them.

Here, I present such a method, developed to systematically compare reptile scalation patterns (often referred to as “pholidosis,”) but applicable in general to surface patterns constructed from units with clear homology relationships. Rather than treating scale numbers and scale adjacency relationships as discrete characters, I draw inspiration from the techniques of geometric morphometrics and anatomical network analysis to analyze the pattern of pholidosis as a network spread across the body of the animal. I describe a simple method for producing these networks and then describe and implement an algorithm that provides an edit-distance measure between networks, allowing for comparative analysis of their topologies–a “topometric” tool. Using a group of fossorial lizards (family Dibamidae) as a study system, I demonstrate the utility of these techniques in understanding the evolution of scalation patterns in a comparative context, investigating the relationships between pholidosis patterns, phylogeny, morphology, ecology and biogeography in this poorly-understood clade. These techniques are implemented in an R package, “pholidosis.”

## INTRODUCTION

Patterns in the bodies of organisms arise through structured development, and the characters that we recognize as differing between species differ because of changes to that developmental pattern. Many characters differ continuously (e.g. length, mass, color), while others differ discretely (count, presence/absence). Some patterns of discrete characters, though, differ between species in complex ways. The scintillating patterns of vein-bordered domains in insect wings and the patterns of scales on the heads of snakes and lizards are made up of discrete units, which in some cases have clear homology relationships between species. These discrete units produce a complex pattern via their presences, absences, and adjacency relationships.

Though these pattens can be distinctive, taxonomically and biomechanically important (Miralles et al., 2011; Salcedo et al., 2019), workers lack a framework by which to compare these topologies at a fine scale. Herpetologists rarely analyze suites of scalation characters, and even studies of insect wing patterning that incorporate network models do so largely to measure coarse network characteristics rather than to investigate fine-scale topological similarity (Hoffmann et al., 2018; Salcedo et al., 2019). To move beyond description and coarse analysis of these patterns into the study of the tempo and mode of their evolution, researchers require tools to directly and consistently compare their topologies.

## PHOLIDOSIS

Squamates are defined by their scales; these keratinous structures sheath their bodies and serve as their main point of contact with their environment. Scales are diverse in color, shape, size, texture, and of course, function (Chang et al., 2009). Studies of the ontogeny (Alibardi, 1997, Tzika et al. 2023), histology (Alibardi, 1997; Dujsebayeva, Ananjeva & Bauer, 2021), and microstructure (Alibardi, 1997; Comanns et al., 2011; Dujsebayeva, Ananjeva & Bauer, 2021) of squamate scales demonstrate how they have been adapted to serve the diverse lifestyles of the squamate radiation, but comparatively little attention has been paid to the inter-relationships of scales on the body surface. This is despite the importance of those inter-relationships to our concepts of squamate species; a discussion of scalation patterns, often termed called “squamation”(e.g. Greer, 1985; Das & Yaakob, 2003), “scutellation”(e.g. Leviton, Brown & Siler 2011) or “pholidosis” (e.g. Scherz et al., 2017; Barros-Filho et al., 2019) is a crucial component in descriptions of most new squamate species (e.g. Da Silva & Ávila-Pires, 2013; Liu et al., 2020; Kliukin et al., 2024b). Many squamate species are diagnosed based on scale counts, sizes, or spatial relationships, but little effort has gone into studying the evolution of these characters; an important oversight given that homoplasy in such relied-on characters could lead to much taxonomic confusion.

As a key characteristic of squamate reptiles, keratinous scales are ubiquitous in lizards and snakes, and are often considered in two separate categories; head scales and body scales. In some taxa, including the majority of snakes, the numbers and spatial arrangements of body scales follow a clear and consistent pattern, organized by the interaction of reaction-diffusion systems and dorsal-ventral positional information (Tzika et al., 2023). Consequently, herpetologists describe patterns of snake body scalation via just a few characters. In some cases, only two characters are necessary; the number of scale rows at mid-body, and the number of ventral scales between the head and the cloaca (e.g. Liu et al., 2020). Conversely, some lizard taxa sport a dazzling array of scale morphologies and interrelationships, with different types scales specialized to their backs, bellies, legs, and tails (e.g. Da Silva & Ávila-Pires, 2013). Head scalation is rarely simple, and the ontogenetic patterning of these scales appears to follow a different logic than body scales (Tzika et al., 2023).

Though heavily relied on for identification, pholidosis patterns are not necessarily fixed for species, and may vary between populations (Gans, 1971; Barros-Filho et al., 2019), between individuals (Gans, 1971; Greer, 1985; Darevsky, 1992; Barros-Filho et al., 2019), or even over the life of an individual (Tomović et al., 2008). In many cases, pholidosis patterns involve both large, placoid scales and small granular scales (e.g. Da Silva & Ávila-Pires, 2013), and the arrangement and homologies of small, granular scales may be impossible to determine. For this study, I focus on large, placoid scales in the study of pholidosis, though the incorporation of fields, patches, or series of scales into a pholidosis network is possible with the methods introduced herein.

### Properties of surface networks

Biological surface networks represent patterns of adjacent units on some surface of an organism. In pholidosis networks, scales are the units, represented in a network by vertices which are connected to adjacent vertices (scales) by edges. Networks of pholidosis are not necessarily unique from other surface networks, but for the sake of simplicity they will be the focus of this paper.

Surface networks have two critical properties:

First, because the units themselves exist on a two-dimensional body surface, surface networks are necessarily planar, meaning that the network can be embedded in a plane such that its edges do not cross (Diestel, 2017). Planarity introduces a constraint into the measurement of edit distances between networks, because it is not possible to arbitrarily link two vertices in a planar network while maintaining its planarity. For instance, the right and left ocular scales of a snake cannot be arbitrarily linked with an edge, because this implies that the right and left eyes have been made adjacent through some untenable folding of the head.

Second, vertices can be considered homologous between networks. In pholidosis networks, this is based on scale homology. Though scale homologies are not universally implied by scale nomenclature in squamates (e.g. Greer, 1985), the assumption of homology between scales or groups of scales in closely-related species of squamates is implied by the consistent use of these characters in taxonomic literature. Just as in other morphological studies, the analyses of surface networks are only as trustworthy as the homology assumptions that underlie them.

Beyond these necessary properties, the networks studied herein are constructed with two additional topological constraints that support the ease of computation of edit distances between networks. First, the networks are undirected, meaning that their edges do not have any directionality; edges simply link pairs of units rather than implying an ordered relationship between them. Second, the networks are fully triangulated (better referred to as maximally planar); that is, that the addition of any edge to the graph would result in a nonplanar graph. Because of this second constraint, the networks can be best thought of as existing on a surface of a sphere rather than on a flat plane, as the networks cannot have any “perimeter.” The network therefore defines the edges and vertices of a polyhedron with all triangular faces. For a regular polyhedron, the number of edges, E, is equal to plus the number of faces, F, Plus the number of vertices, V, plus 2 (E = F+V+2). In a triangulated polyhedron, there are tree faces for every two edges (E = 3/2 F) (Bowen & Fisk, 1967). Therefore, the networks described herein have a simple relationship between the number of edges and the number of faces: E = 3V – 6.

Finally, there are some inter-network relationships that the methodology described herein cannot account for. In the method of measuring network edit distance used herein, large changes in the position of some subset of scales relative to another subset are equivalent to the complete separation and re-attachment of these sub-networks (Supplementary figure S1). Step-wise movement of the subsets relative to each other is not explicitly modeled via the algorithm presented here, as no such movement is required to calculate edit distances among the exemplar dataset of dibamid lizards.

The properties of squamate scales can be embedded in these networks as attributes of edges or vertices. In this study, I demonstrate the use of edge weight to denote fusion or near-fusion of scales, but in principle scalation networks can embed many more types of data; for edges, the length of the shared scale perimeter or the presence and directionality of overlap; for scales, size, shape, color, surface texture, ossification, innervation, special sensory or exocrine function, etc. The challenge of analysis of any character lies in implementing a reasonable model of character state transition. In the methods described and implemented herein, only topological characters (presence, adjacency, and fusions) are considered.

### Dibamid lizards (family Dibamidae)

Lizards of the family Dibamidae are among the most poorly understood reptiles on Earth. These wormlike lizards have highly reduced eyes covered by an ocular scale, no forelimbs, and flap-like hindlimbs only in males. They are presumed to be fossorial insectivores, though precious little has been published on their diet or behavior (see Darevsky, 1992). Though dibamids resemble amphisbaenians and fossorial skinks, they are not closely related to either group; recent phylogenomic studies place Dibamidae as sister either to Gekkota (Burbrink et al., 2020) or to all other squamates (Singhal et al., 2021). Twenty-seven species of *Dibamus* live in the tropics of Southeast Asia, Indonesia, the Philippines, and New Guinea (Kliukin et al., 2024a), while *Anelytropsis papillosus* is found in Mexico. Despite this major geographic rift, *Anelytropsis* appears to be most closely related to a clade of *Dibamus* residing in mainland Southeast Asia. This clade is sister to a “Peninsular-Island” clade of *Dibamus* inhabiting Peninsular Malaysia, Indonesia, the Philippines, and New Guinea (Townsend, Leavitt & Reeder, 2011).

Dibamids are quite conservative in their external morphology (Greer, 1985; Quah et al., 2017), differing primarily in scalation patterns, body size, and tail length, and no morphological characters are known that distinguish the *Dibamus* clades recovered by Townsend, Leavitt & Reeder (2011). Most dibamids exhibit a “rostral shield,” of scales fused into a single keratinous structure covering the snout, though the structure is quite variable; different species exhibit many different phenotypes of scale fusion.

From the available data, dibamid pholidosis characters are largely stable at the species level (Greer, 1985; Quah et al., 2017), though a lack of known specimens may contribute to an underestimation of variability. There is known intraspecific variation in the number of post-ocular and post-infralabial scales in some species, including the apparently quite variable and widespread *D. taylori* (Greer, 1985), and variability in the degree of fusion between scales of the rostral shield in some species (e.g., *Dibamus bourreti*; see Greer, 1985; Darevsky, 1992). As this study is primarily a demonstration of pholidosis network methodologies, rather than a particularly in-depth study of dibamid evolution, I represent each dibamid species using only a single scale configuration.

## METHODS – PHOLIDOSIS NETWORKS

**Figure 1.**
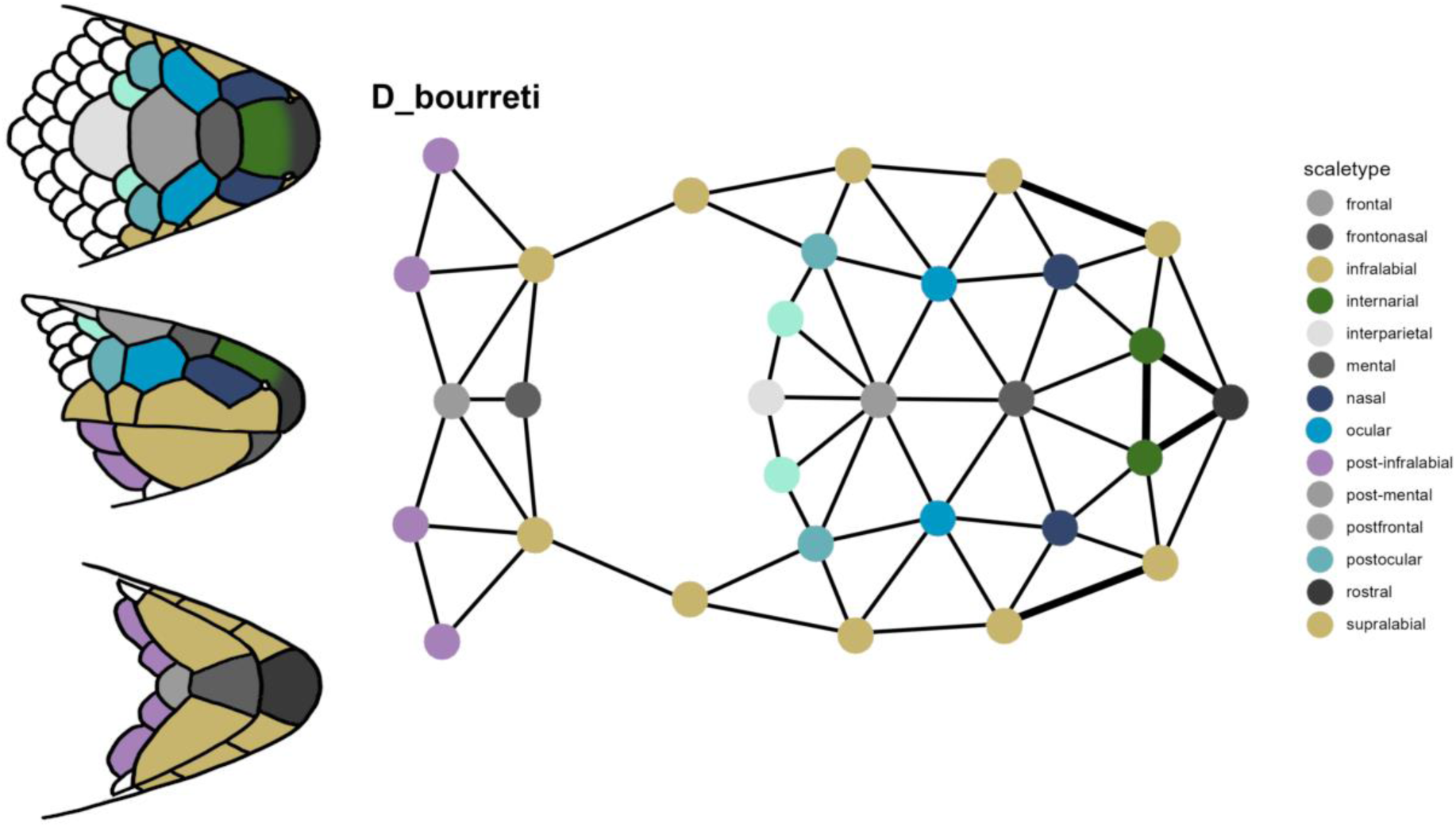
Pholidosis network of Dibamus bourreit. A. Dorsal, lateral, and ventral views of Dibamus bourreti (redrawn after Darevsky (1992) but with the fused supralabials reported by Greer (1985)) with scales colored according to their scale type. B. The resulting pholidosis network; thick lines indicate fusions between scales.

All of the methods described herein are implemented in an R package, *pholidosis*. A brief guide to using the package can be found in an accompanying publication (Krone, 2025). The package can be downloaded at github.com/dibamus/pholidosis.

### Encoding the network using an adjacency matrix

The first step in quantifying surface pattern is to encode it in a machine-readable format. I follow (Esteve-Altava & Rasskin-Gutman, 2014) in beginning with an adjacency matrix, encoded as follows:

Let *S* be the set of scales involved in a pholidosis pattern. *S* contains *n* scales (*s_1_, s_2_, … s_n_*). *M* is a square matrix of dimensions n*n, where each scale in S occupies a row and column in this matrix. If a scale *s_i_* borders a scale *s_j_*, the matrix cell corresponding to [s_i_,s_j_] is filled, producing an edge between those vertices. If the scales are adjacent and fully separated, I denote this with the number 1. Scales that are partially fused together are denoted by the number 2, while scales that are fully fused are denoted by the number 3. On importation into R, this number encodes an “edge weight” in the resulting network, *G*.

I find the best practice when creating these matrices is to include in *S* all possible scales within the pholidosis patterns of all species involved, as well as any structures that these scales abut, such as the eyes, mouth, and remaining body surface. If some pholidosis patterns have two pretemporal scales (*s_pretemporal1_, s_pretemporal2_*), while others have only one (*s_pretemporal1_*), the same blank matrix *M*, containing rows and columns for both supratemporal scales, can be filled in to describe the whole set of patterns; matrices corresponding to species lacking the second pretemporal scale will simply have a corresponding blank row and column. This blank row and column produces an “isolated” vertex in the network, which can be removed for analysis.

The *pholidosis* package uses *igraph* (Csárdi & Nepusz, 2006) and the *igraph* R package (Csárdi et al., 2025) to represent and perform operations on networks.

### Network edit distance

Surface networks can be compared by a measurement of edit distance, the number of changes needed to change one network into another. Any network can be transformed into any other network by the addition and subtraction of edges (and therefore vertices), but simply counting the number of different edges between two networks can produce an overestimate of the number of biological changes needed to change one network into another. For instance, the addition or subtraction of a scale from a pholidosis network should be considered a single biological change, but this can involve the addition or subtraction of any number of edges, depending on what scales the new scale is adjacent to. Likewise, some shifts in the adjacency relationships of scales shared by both networks involve both the addition and subtraction of edges (Figure 2), though a biological interpretation implies a simpler process, in which a single change, for instance, the enlargement of a scale, necessitates cascading effects.

**Figure 2.**
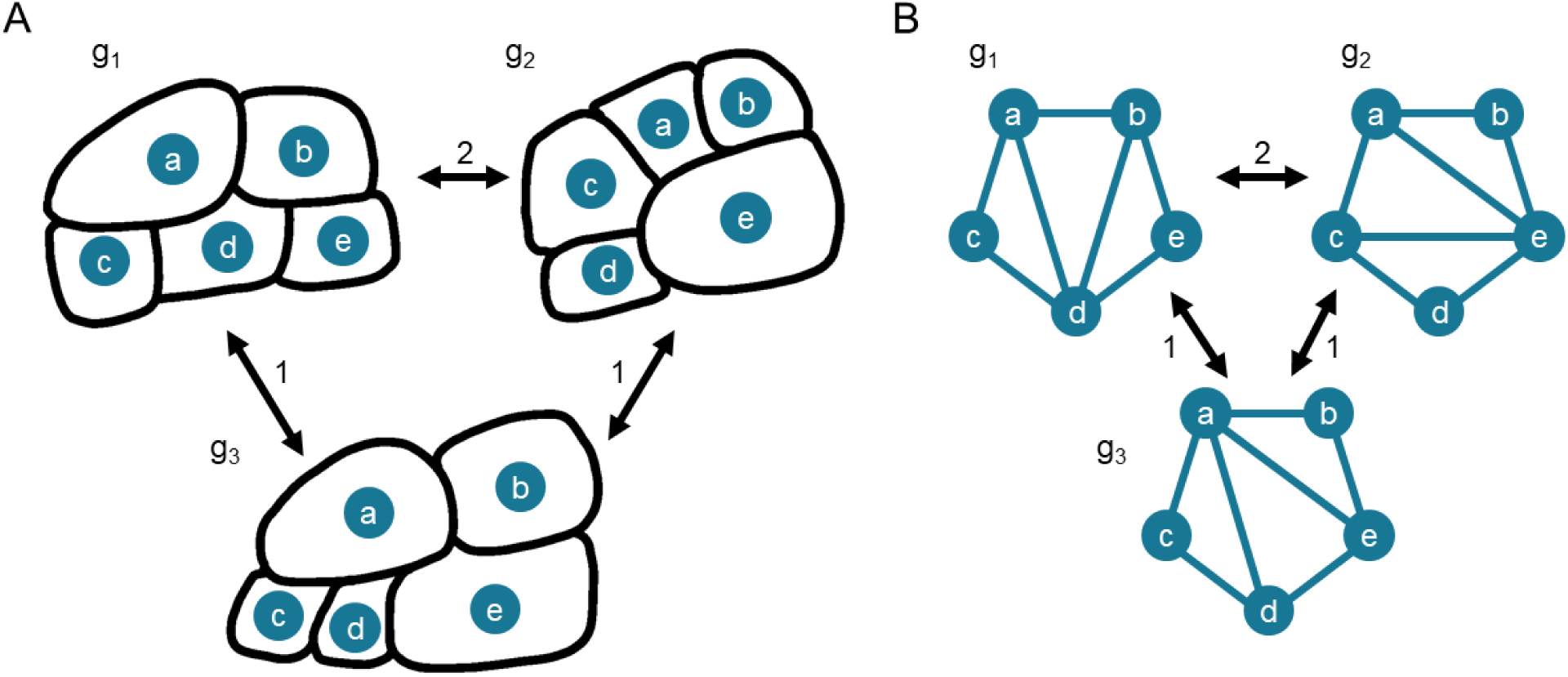
– Edit distance accounting for topological changes. A. Three pholidosis networks (g_1_, g_2_, and g_3_) involving five scales, a, b, c, d, & e B. The same pholidosis networks represented as networks rather than physical arrangements of scales, with adjacency relationships between the scales represented by edges. The five vertices, a, b, c, d, & e, form a cycle with n-3 = 2 internal edges. Note that the full graphs also contain sub-cycles of four vertices with a single internal edge. Arrows between the networks track how many changes it takes to transform one network into the other (the edit distance). Though the networks g_1_, g_2_, and g_3_ all contain the same scales/nodes, their topologies are different. To transform g_1_ into g_3_, the edge (b, d) is removed and the edge (a, e) added. We can see that this topological change is the result of scale e growing in size such that it contacts scale a, therefore necessitating the loss of a contact between b and d. Therefore, though this change involves the loss of one edge and the gain of one edge, it is counted as a single change, since the gain and loss cannot occur separately in this arrangement, as the intermediate state would violate the need for the network to be fully triangulated. By a similar process, g_3_ can be transformed into g_2_; this is the “edge swap.” Notice that from g_1_ to g_2_, there is a similar loss of two edges (a, d) and (b, d) and a gain of two edges (a, e) and (c, e). The transformation of g_1_ into g_2_ may occur through an intermediate scale arrangement (g_3_), but this intermediate step is not modeled, as we can assume that two changes have occurred between g_1_ and g_2_, since the two 5-vertex cycles of nodes surround unmatched edges, of which there are necessarily 2.

The following method is sufficient to account for these cases in the dibamid dataset, producing an estimate for the minimum number of changes in vertices, edges, and edge weights to transform one scalation network into another. It is implemented in the *pholidosis* package by the *pnet_distance* and *graph_edit_distance* functions.

The *graph_edit_distance* function measures the number of topological changes needed to transform one network, *G*, into a second network, *H*.

In order to tabulate and keep track of the necessary changes to transform one graph into another, we must create descriptive tables of edge properties for the graphs *G* and *H.* For each edge in *G*, we assess the following properties: 1) the names of the vertices that the edge connects; 2) matched edges: whether an edge in *H* connects two vertices with the same names; 3) alternate edge paths: if no edge in *H* connects vertices of the same names, the shortest path (if any exists) between two vertices with the same names in *H* via edges unique to *H*; 4) whether, in those alternate edge paths, all intermediate vertices are unique to *H*; and 5), whether the edge is “resolved” (can be linked to an identical or alternative edge in *H*). We produce the same table for *H* with respect to *G*. The *pholidosis graph_edit_distance* function produces this table as an R data frame, and relies on the *igraph* function *shortest_paths* to find alternative paths between vertices.

If an edge in *G* 1) does not contain unique vertices, 2) is not matched by an edge in *H*, and 3) its vertices cannot be connected by a path in *H* that contains only vertices unique to *H*, it is considered “unresolved”, and marked as such in the descriptive table. These unresolved edges represent topological differences between *G* and *H* that involve only vertices found in both networks (Figure 2). In order to determine the smallest number of changes required to transform *G* into *H*, we must determine if these unresolved edges represent cases of edge “swapping,” in which sub-networks of *G* and *H* share a ring of homologous vertices surrounding different internal edges (Figure 2B). In these “swapping” cases, simply comparing the number of unshared edges between *G* and *H* would double-count simple changes in scalation (Figure 2).

When descriptive tables for the edges of *G* and *H* have been produced, we compare their unresolved edges to identify these systems of “swapping” and count the number of edges in *G* involved in the systems. This is achieved in a few steps. First, for all unresolved edges, we identify the two triangles (3-cycles) that the missing edge was a part of; since each graph is fully triangulated, every edge of the graph is part of exactly two triangles. One can visualize this as a square with a diagonal edge from corner to corner; that diagonal is the unresolved edge, and we can call this 4-vertex subgraph the unresolved edge’s “square graph”. In order to define the two 3-cycles, we must know the names of the vertices in the square graph that are not linked by the unresolved edge. Once these two vertices of the square graph are found for each unresolved edge, we construct an adjacency matrix of unresolved edges; unresolved edges are adjacent if they both appear in each others’ square graphs.

In order to identify corresponding systems of unresolved edges in *G* and *H*, we must identify systems of unresolved edges linked by this adjacency (e.g., internal edges in Figure 2B). From this adjacency matrix, we construct a summary graph wherein the identified unresolved edges are vertices, and the adjacent vertices are linked by edges. We then identify connected vertices, here corresponding to distinct groups of “problem edges” linked by adjacency. We then assign the problem edges to these groups. These groups are then identified by the set of vertices in the original graph that are part of the unresolved edges and their square graphs; this set of vertices forms a cycle around the unresolved edges of length equal to the number of unique vertices of the unresolved edges plus two vertices from the square graphs that were not involved in adjacent edges. The names of vertices in this cycle are then added to the descriptive table. In the *pholidosis* function *graph_edit_distance*, identification of the unresolved edge groups is achieved via a depth-first search of the summary graph using the *igraph* function *dfs*.

We then compare these cycles between graphs G and H. If both the cycles and number of unresolved edges in each cycle are identical between the graphs, all of these unresolved edges are considered “resolved.” In this case, the number of edge-swaps to turn *G* into *H* is equal to the number of unresolved edges in *G* or *H*. If there is any difference between the recovered cycles and number of edges in the cycles between the graphs, then our methodology has failed; the differences between *G* and *H* are too great for this method to resolve an unambiguous edit distance (see Supplementary Figure S1). In this case, the *graph_edit_distance* function returns an NA value for the number of edge differences.

With descriptive tables for *G* and *H*, we can count changes in the number of vertives and changes in the structure of edges in *G* and *H* (node changes). We call these the *f* (vertex) and *e* (edge) distances. In graphs with differing edge weights denoting fusion or partial fusion of vertex elements, we can also count the number of changes in edge weight differences between the networks, the *w* distance. Since the fusion of two scales should be evolutionarily equivalent to the deletion of one scale, full fusions are counted as 1 change, and semi-fusions as 0.5 changes.

The *pholidosis* function *pnet_distance* produces an edit distance between two networks using *graph_edit_distance* and counts these weight differences, returning three distance measures; *f*, the number of vertex differences between the networks; *e*, the number of edge differences between the networks, and *w*, the number of edge weight differences between the networks. If desired, it will also return the descriptive tables for *G* and *H* as data frames produced by *graph_edit_distance*.

In some cases, the algorithm outlined here is insufficient to resolve the edit distance. Some of these involve real large shifts in the position of some graph elements relative to others; an example of this is given in supplementary figure S1. In other cases, insufficiency may be due to an incorrect homology assumption that produces a drastic shift in the network. For instance, in several *Dibamus* species such as *D. seramensis*, a scale located behind the ocular scale separates the ocular and postfrontal scales. In the case where this scale is labelled “postocular 1,” and the species has a series of 4 postoculars, the *graph_edit_distance* function fails to calculate a distance between species with and without this scale arrangement. However, when we consider this a new type of scale, a “parafrontal,” our algorithm can calculate distances between animals with and without this scale arrangement.

### Distance and character matrices for pholidosis networks

The *pholidosis* package contains functions to build both distance and character matrices from collections of pholidosis networks. Given a list of pholidosis networks, the function *net_dist_mat* calculates distance measures *f*, *e*, and *w* for each possible pair of networks in the list using *graph_edit_distance*, and returns a 3*n*n array, containing matrices for *f*, *e*, and *w* distances between n networks. This distance matrix is a measure of distances between pholidosis networks in topospace, analogous to the morphospace in a geometric morphometric analysis.

Pholidosis networks can also be broken down into sets of atomized traits. The *pholidosis* function *morpho_matrix* produces a morphological character matrix that treats edges in a network as individual characters. The matrix rows correspond to the pholidosis networks, and the matrix contains a column for each unique edge in the list of networks. If a given edge is present in a given network, the corresponding cell in the matrix is scored with that edge’s weight.

### Network edit distance as a metric

In order for the network edit distance to be a useful measure, it should define a metric space. A metric space is a space in which four conditions hold; 1) that the distance from a point to itself is zero; 2) that the distance between any two distinct points is positive; 3) that the distance from point *a* to point *b* is the same as point *b* to point *a*; and 4) that the triangle inequality holds true. The triangle inequality states that the distance between points *a* and *b* must be less than or equal to the sum of the distances from *a* to *c* and *b* to *c*.

The first two of these conditions are necessarily fulfilled by the network edit distance calculated herein; all differences between networks are accounted for in the course of measurement by the descriptive tables and each difference is assigned some positive value. Since networks are compared pairwise and neither is given any preference by the edit distance algorithm, the third condition holds.

Investigating the condition of the triangle inequality is more complex. Since the *f* distance is based on counts of unshared vertices, this intuitively must be true. All differences are measured in positive single units, so any vertices unique to graph *C* will increase its distance from graphs *A* and *B*. For the *e* distance, we assume axiomatically that the smallest change between graphs is 1 edge swap. This type of change, a swap of the central “chord” edge in a 4-cycle (figure 2B, g_1_ to g_3_), provides the basis of the metric. Figure 2B demonstrates the equivalency of the swapping of two chords in a 5-cycle to two steps of swapping in 4-cycles. This logic can be extended to larger cycles with more chords. One can find a series of changes such that each edge in *A* can be swapped for an edge in *B* to resolve one square graph at a time in this larger cycle. This produces the result that the number of swaps required to change *A* into *B* is equal to the number of edges in *A* that are not found in *B*, so long as all of those edges are chords of homologous cycles of *A* and *B*. As before, edges unique to graph *C* will only increase its distance from graphs *A* and *B*.

We can check whether the implementation of this graph edit distance algorithm in *pholidosis* produces any triads of networks in the dataset that violate the triangle inequality. For the dibamid dataset here, the inequality holds.

## METHODS – PHOLIDOSIS NETWORKS OF DIBAMID LIZARDS (DIBAMIDAE)

All code and input data needed to replicate these analyses can be found at

I referenced diagrams and photographs from literature sources to code scale adjacency matrices for all published dibamid species, basing the matrices on the holotypes whenever possible (Supplementary Materials: Appendix). I am grateful for the help of Dr. S.R. Chandramouli who sent me high-resolution photos of *Dibamus nicobaricum*, without which I could not have fully resolved its pholidosis pattern (personal communication). The scalation patterns in *D. montanus* and *A. papillosus* elucidated which scales fused together to make the rostral shield in other species. These include the rostral scale, the first three supralabial scales, right and left nasal scales, and right and left internarial scales. In the adjacency matrices, adjacent scales were coded as adjacent (1), partially fused (2), or fully fused (3), but these values are changed to adjacent (1), partially fused (1.5), or fully fused (2) on importation via a call to a setup function, by default, *lizard_setup*, by the *scale_network* function.

To calculate a distance matrix for the set of 27 dibamid species (all except the recently published *Dibamus elephantinus*), I used the *net_dist_mat* function implemented in *pholidosis*. The three components of the distance matrix (the *w*, *e*, and *f* distances) were added together to obtain a single distance measurement between each species pair. I then performed Principal Coordinate Analysis on the summed dibamid distance matrix to map dibamid scale network diversity onto two major axes of variation using the base R function *cmdscale*. I used the *WDBdisc* function in the R package *WeDiBaDis* (Irigoien, Mestres & Arenas, 2017) to perform a unweighted discriminant function analysis based on a reduced version of this distance matrix (excluding *Anelytropsis papillosus*) to attempt to distinguish between Mainland and Peninsular/island *Dibamus* groups.

Maximum snout-vent length (SVL) and maximum proportional tail length measures used in this paper were compiled by Quah et al. (2017) and supplemented with data from Koppetsch, Böhme & Koch, (2019), and Kliukin et al. (2023,2024).

Since no comprehensive phylogenetic hypothesis exists for Dibamidae, dibamid scalation cannot be tested within a phylogenetic context. To account for some degree of possible phylogenetic signal in scalation patterns, I categorized species into three geographic groups (Mainland Southeast Asia, Peninsular & Island Southeast Asia, and Mexico) following the clades identified by Townsend, Leavitt & Reeder (2011).

From the scale networks, I calculated the number of scales, number of fusion events, network strength, and composite scale count (scale count in which fully fused scales are counted as a single scale). I built linear models with the base R *lm* function using these summary statistics, with and without location as a second term, to understand the basis of the principal axes identified by the principal coordinates analyses. I then compared these eight models for both axes in an AIC framework using the *aictab* function of the *AICcmodavg* package (Mazerolle, 2023).

Using the same approach, I investigated covariation between head scalation and body shape (SVL and maximum proportional tail length). I built linear models using the network summary statistics and Principal Coordinates as predictors, with and without location as a second term. Again, I compared these twelve models for each body shape variable within an AIC framework.

I used distance- and parsimony-based methods to generate phylogenetic hypotheses for Dibamidae based on pholidosis network data alone. From the pholidosis network distance matrix, I produced a distance-based tree following the algorithm by Saitou & Nei (1987) using the *ape* package’s neighbor-joining function *nj* (Paradis & Schliep, 2019). For the parsimony tree, the scalation networks were decomposed into discrete characters (one for each edge) using the *pholidosis morpho_matrix* function. I then performed a maximum parsimony analysis of this matrix, with edge weight coded as an ordered state, using the phangorn function *pratchet* (Schliep, 2011).

## RESULTS

### Dibamus networks

Edit distances between dibamid species (summing differences in scale count, scale fusion, and topological differences between scale configurations) are normally distributed (Shapiro-Wilks W = 0.99, p = 0.003) Edit distances range from 1 (*D. tropcentr* to *D. smithi*) to 26 (*D. bourreti* to *D. manadotuaensis*), with a median of 11 and a mean of 10.93. Across the distance matrix, edge weight differences accounted for a total of 1727 steps of edit distance, edge topology changes 102 steps, and additions and subtractions of vertices for 2008 steps.

Dimensionality reduction via principal coordinates analysis of the network distance matrix into two dimensions fails to separate dibamids geographically (Figure 3). K-means clustering of principal coordinate data with k=2 and k = 3 groups does not identify geographically coherent sets of taxa; “Mainland” and “Peninsular-Island” groups occupy overlapping and approximately equal areas of morphospace. With k=2 groups, the species are partitioned primarily along PCoA 1; first a small group with PCoA 1 values of five and above containing *Anelytropsis papillosus*, *D. bogadeki, D. bourreti, D. deharvengi, D. greeri* and *D. somsaki*, plus *D. montanus* with PCoA 1 value of just over 2; and second, a larger group at PCoA 1 values below 2.5 containing all other Dibamids. Correspondingly, distance-based discriminant function analysis could not fully distinguish between Mainland and Peninsular/island groups of Dibamus, finding a classifier function that grouped two mainland *Dibamus* species with the Peninsular/island group.

**Figure 3.**
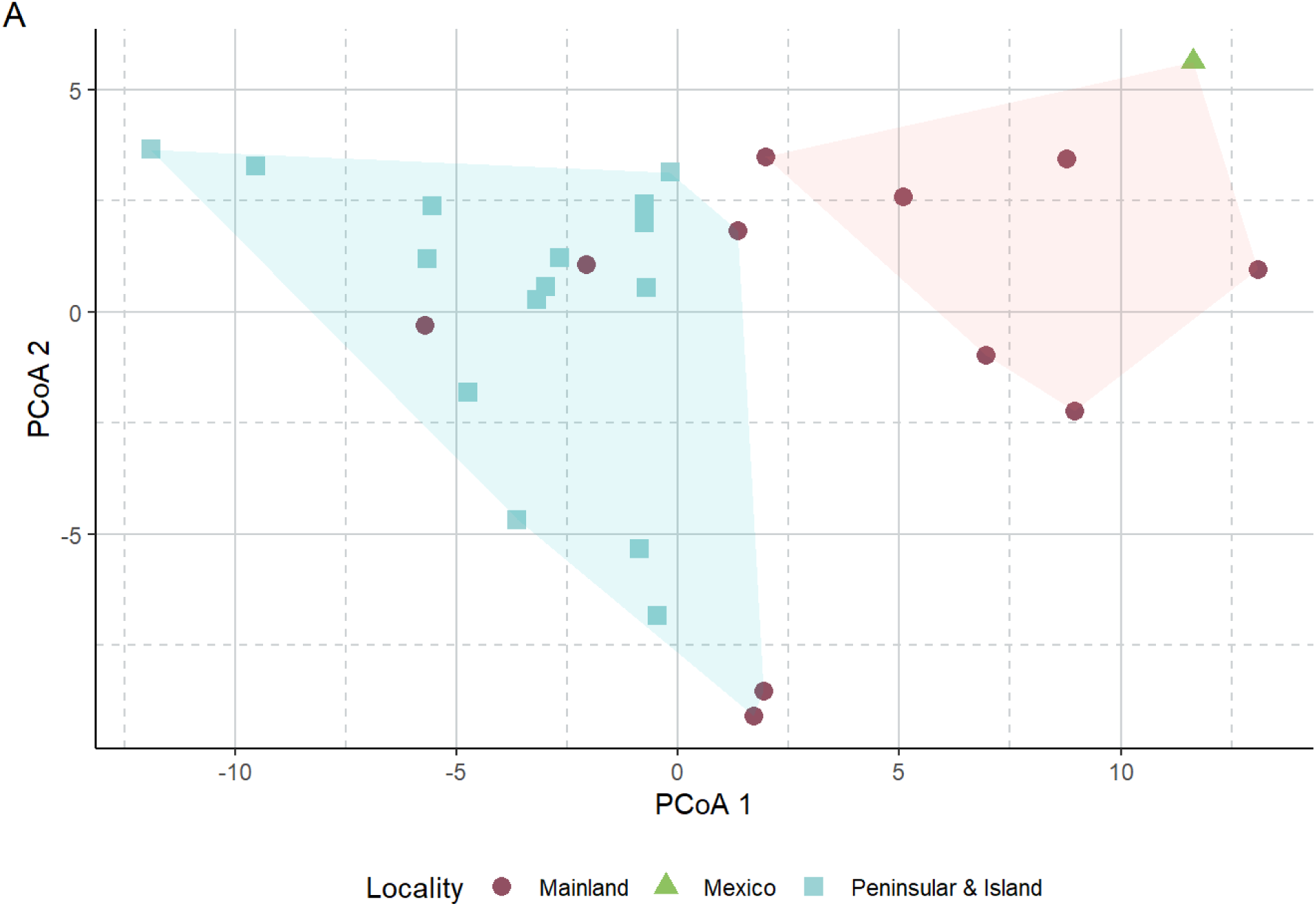
– Dibamid scalation topospace. This principal coordinates plot reduces dibamid head scalation network topospace to two dimensions. Blue squares represent Peninsular/island species while purple circles represent Mainland species. Anelytropsis papillosus is represented by a green triangle. Shaded areas identify clusters identified with K-means clustering (k=2).

The first principal coordinate axis is best explained by network strength, while PCoA 2 is best predicted by the number of fusion events and locality (Figure 4). The number of fusion events and locality jointly explain 82% of the variation in this axis (Supplementary Table S1).

**Figure 4.**
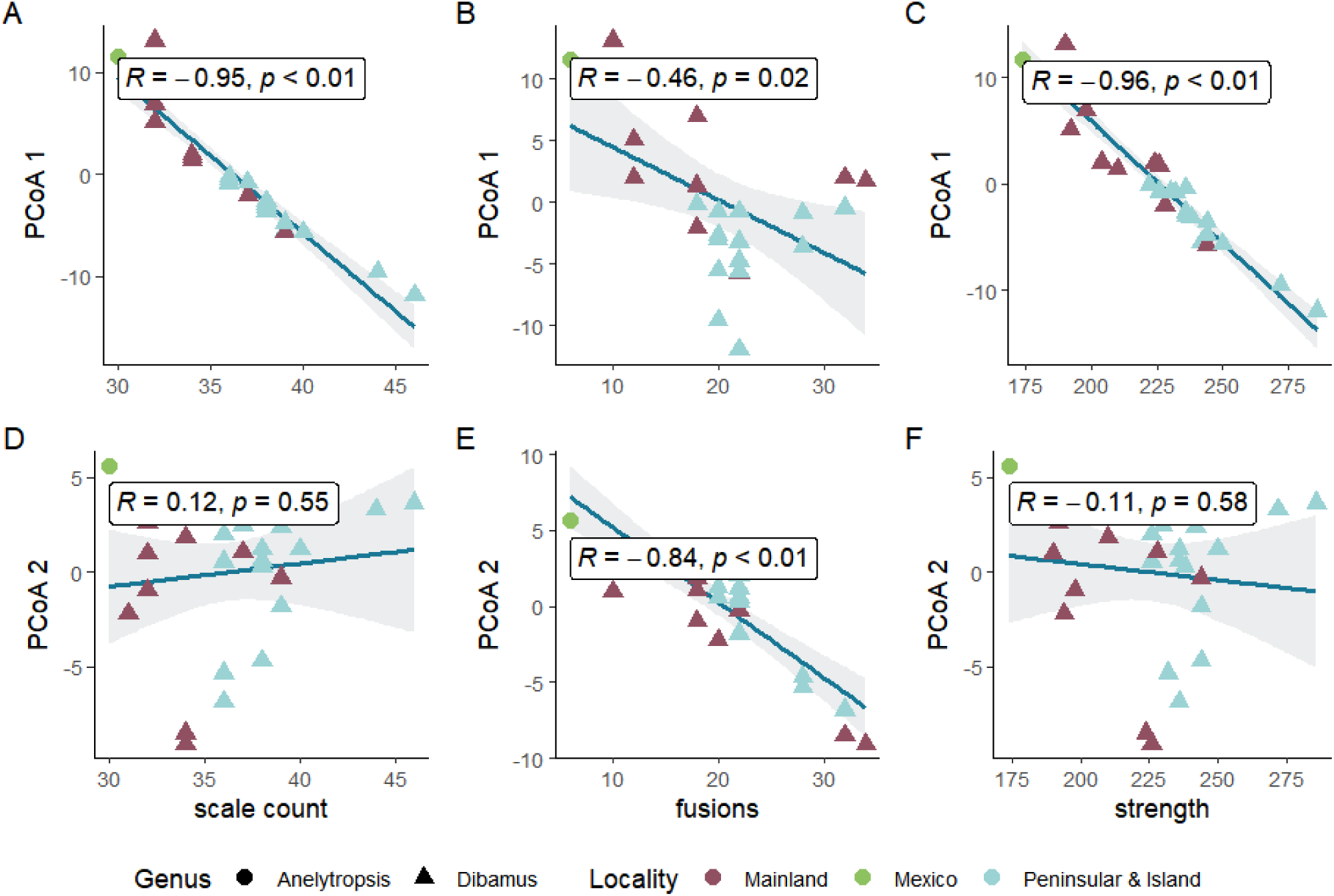
– Correaltions of principal Coordinate axes (PCoA) and graph properties.

Dibamid pholidosis networks are influenced by body size (SVL). Maximum SVL is correlated with PCoA 2 (R^2^=0.55, p< 0.01) and with the number of fusion events, with larger species having fewer fusions (R^2^=-0.41, p = 0.03) (Figure 5D).

**Figure 5.**
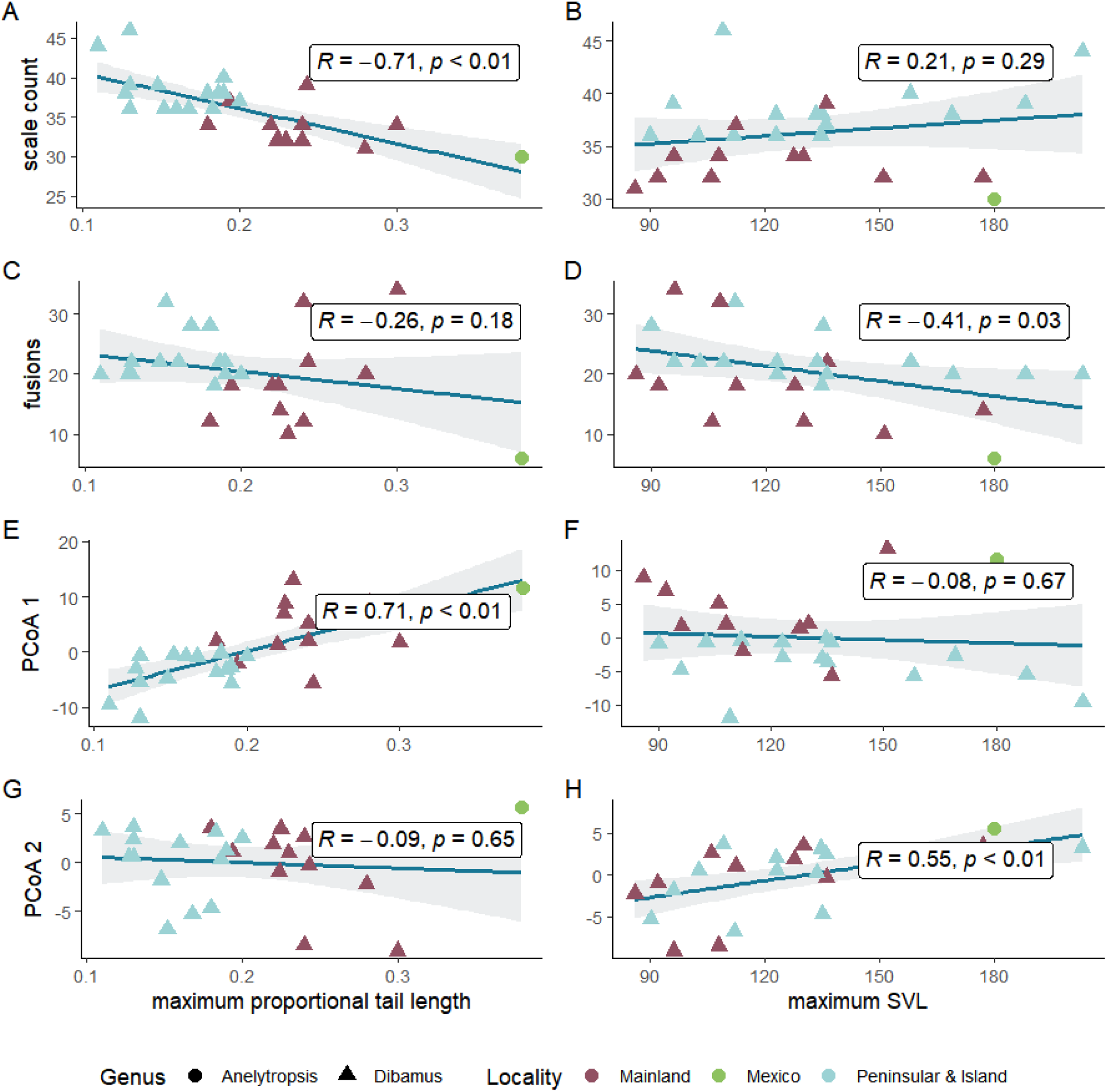
– Regressions of dibamid head scalation network properties (vertical axes) on tail length and snout-vent length (SVL) (horizontal axes)

Surprisingly, pholidosis patterns of dibamid heads are strongly connected to tail length. Tail length is a probable proxy for fossoriality (Wiens, 2006; Lawing, Head & Polly, 2012) that is not correlated with SVL in dibamids (R = −.03, p = 0.84). Figure 5 (A,C,E) shows that scale count and therefore PCoA 1 are tightly correlated with maximum proportional tail length (both R^2^>0.73, all p<0.01). However, Mainland and Peninsular/island *Dibamus* groups have very different average tail lengths; the mainland species tend to have much longer tails (Welch’s t = −5.9, df = 19.4, p < 0.001) and correspondingly weaker/smaller scalation networks (Welch’s t = 4.5, df = 21.0, p < 0.001). Models including locality and either PCoA 2 values or composite scale count best explained differences in tail length in dibamids, with locality the major predictor (Supplementary Table S2).

Scalation characters do not resolve particularly robust phylogenies for Dibamidae. The consensus parsimony tree includes several polytomies and has a consistency index of 0.52 and retention index of 0.72, indicating high homoplasy. The neighbor-joining tree

Both distance- and parsimony-based phylogenetic hypotheses agree partially with the Bayesian phylogeny of Townsend, Leavitt & Reeder (2011) (Figure 6). Most saliently, they embed *Anelyropsis papillosus* within a clade containing the “mainland” species, rendering that “mainland” clade paraphyletic. Both trees also recover a primarily Sulawesi-adjacent clade (*D. celebensis*, *D. manadotuaensis*, and *D. seramensis*) rendered paraphyletic by *D. deimontis* from Vietnam. The neighbor-joining tree recovers the “Peninsular/Island” clade with the same branching order as the molecular tree; if re-rooted such that the “Mainland” clade would be sister to all other dibamids, the parsimony tree would recover the same.

**Figure 6.**
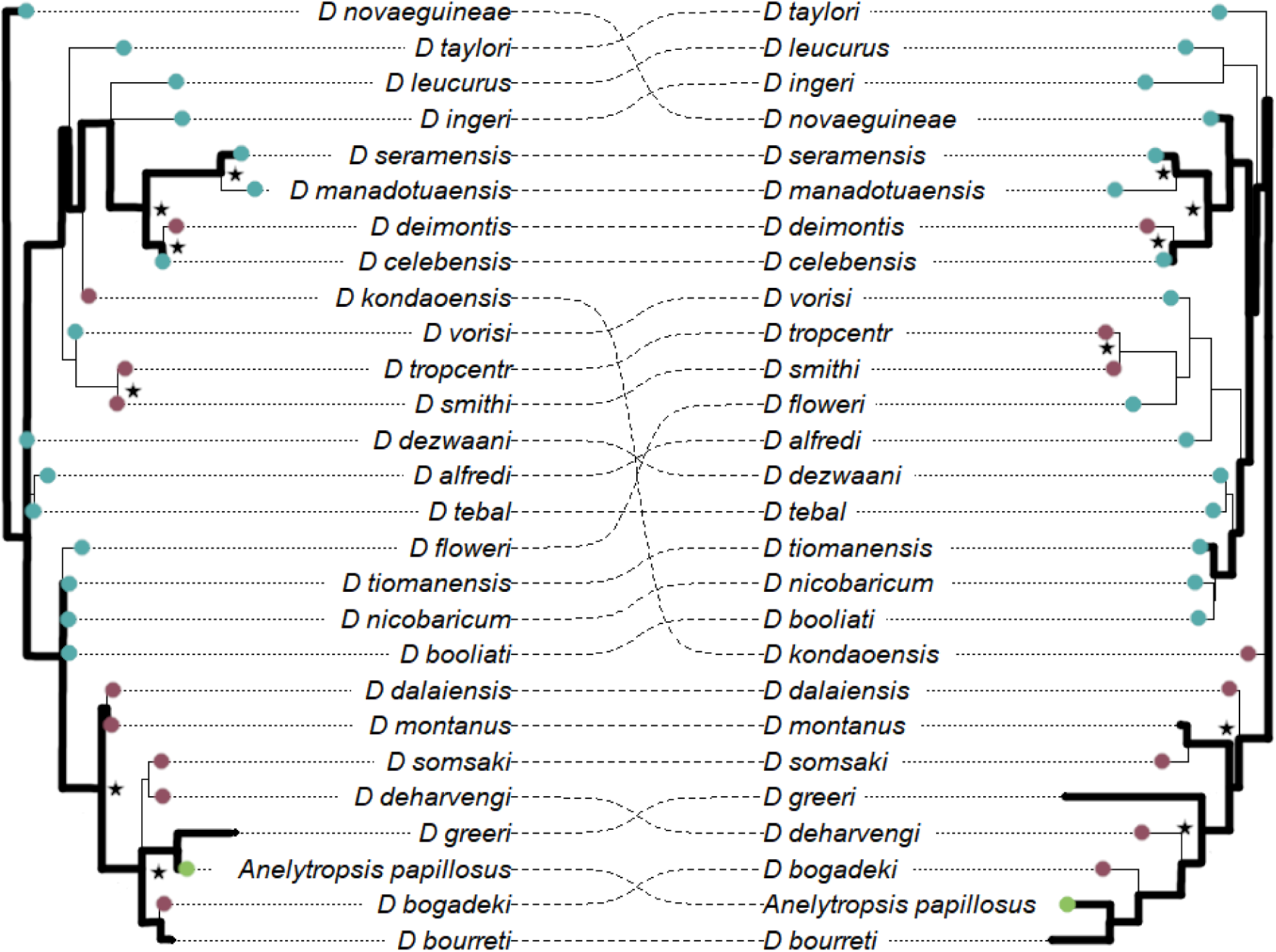
– Phylogenies for Dibamidae based on scalation networks. The left tree is a consensus maximum parsimony tree based on a matrix of edge states for all dibamid pholidosis networks. The right tree is a neighbor-joining tree based on graph edit distances between those networks. Mainland species are indicated with purple, peninsular/island species are indicated with blue. Bold edges connect species found in Townsend, Leavitt & Reeder (2011). Clades found in both trees are marked with a star.

However, the phylogenies have rather low agreement; few clades near the tips of the trees are shared between the two, and quartet agreement between the two trees is 52% (9165 agreeing quartets of 17550 possible quartets).

## DISCUSSION

### Using surface networks

By connecting a wealth of otherwise atomized character states into a biologically representative mathematical structure, the methods described herein allow for rigorous comparative analysis of surface patterns. Here, they have produced insight into possible allometric and ecological patterns in a secretive and poorly-understood group of lizards. The distance measurement method described here produces biologically intuitive results when comparing networks using principal coordinates analysis, discriminant function analysis, and tree estimation techniques.

### Dibamid evolution and ecology

The evolution of dibamid pholidosis networks appears to be slow and piece-wise, yet still related to the animals’ ecology. Peninsular/island and mainland *Dibamus* species do not appear to be differentiable based on any single character despite a putative crown age of 54–95 million years for Dibamidae (Townsend, Leavitt & Reeder, 2011). Moreover, though scalation networks produce poorly-resolved phylogenetic hypotheses, they still do appear to carry some amount of phylogenetic information. This suggests an extremely slow rate of evolution in scale characters, as a high rate of character state changes among the small number of characters involved would likely overprint this signal over such a long timespan.

It is surprising that scalation patterns seem capable of resolving biogeographic structure within *Dibamus*. While the phylogenetic hypotheses produced by scalation data are far from robust, they do suggest a more complex phylogenetic structure than a simple Mainland/ Peninsular & Island split. Despite living in Vietnam, *Dibamus kondaoensis*, *D. deimontis*, *D. smithi*, and *D. tropcentr* all appear outside of the “Mainland clade” of *Dibamus*. Nevertheless, the differentiability of Mainland and Peninsular/Island Dibamus based on tail length suggests that Townsend et al.’s clades are ecologically distinct, with the short-tailed Peninsular/Island *Dibamus* species being more obligately endogeic.

Pholidosis patterns may be adaptive in Dibamidae. However, if shorter tails are indicative of more fossorial lifestyles, the number of fusion events (and therefore the degree of development of the “rostral shield”) does not necessarily carry the same signal. Given the apparent morphological split between Mainland and Peninsular/Island *Dibamus*, it may be that the development of the “rostral shield” is merely overprinted in dibamids by a strong phylogenetic signal. Alternatively, rostral shield development may be a consequence of miniaturization more than adaptation, as body size and number of fusion events in dibamid pholidosis networks are inversely correlated (Figure 5). Even with a comprehensive dibamid phylogeny in hand, the adaptive significance of these scalation patterns will remain unclear until these obscure lizards’ ecology is much better known.

Intraspecific variability in pholidosis may complicate these conclusions. The differences between some very similar *Dibamus* species are attributable to a single change in scale count or single incomplete fusion event, and intraspecific variation is known to exceed this in many species (see Kliukin et al., 2024). Future taxonomic work on dibamids informed by molecular phylogenetics may re-partition some of this intraspecific variation into interspecific variation. Until more dibamid specimens are carefully examined and correctly assigned to species, the degree to which scalation patterns are inter- and intraspecifically variable will be unclear. But the relationship between body size and scalation on the interspecific level, combined with the known variability within populations, suggests that there may be an ontogenetic component to dibamid pholidosis, which my examinations of some series of *Dibamus* appear to support.

### Future directions in topometrics

I hope that herpetologists will use the tools presented here to rigorously investigate pholidosis, a long-overlooked aspect of squamate evolution. These preliminary results are encouraging; pholidosis patterns appear to retain some phylogenetic information and appear to be influenced by allometry and ecology. But dibamids are likely a poor model of general pholidosis evolution, just as they would be a poor model for general squamate morphology or ecology. Expansion of these methods to diverse groups of reptiles will certainly provide a better understanding of the evolution of pholidosis patterns in squamates and likely result in better-informed methods of species identification.

By providing comparative tools for the study of biological surface networks, I hope to encourage others to take an interest in these patterns and attempt their comparative analysis. This methodology is ripe for further refinement and expansion. The distance metric implemented in this study will likely be sufficient to analyze a large number of surface patterns, but could be expanded to process large-scale shifts in network structure that might occur in other organisms, such as between families or orders of insects (Salcedo et al., 2019). Further incorporation of vertex- and edge-specific characters and models of their transition could prove useful in studying aspects of surface networks beyond bare topology, including surface properties such as texture, shape, and color. A fully developed methodology of topometric comparison and vertex characters, including edge shape and surface color could, for instance, be used to completely describe the appearance of a butterfly wing, and to quantify the interplay between methods of reaction-diffusion and positional information in producing differing color patterns between different species of butterflies.

## Supporting information

Code for analysis

## ACKNOWLEDGEMENTS

I am indebted to Drs. Marvalee H. Wake and Simon G. Scarpetta for their insightful comments on early drafts of this paper, and to Dr. S. R. Chandaramouli for sharing images of *D. nicobaricum*.

