## Supplementary material for "A network-based comparative method to study reptile scalation and other homologous two-dimensional patterns": Code for analysis: Dibamus_scale_networks.Rmd

````
---
title: "Dibamid Pholidosis"
author: "--"
date: "2025-02-09"
output: html_document
---
# Setup
## Install and load the pholidosis package

Install the 'pholidosis' package. This requires devtools.

```{r install }
library('devtools')
library('remotes')
install_github("dibamus/pholidosis")
```

```{r load packages}
library('pholidosis')
```

# Use *pholidosis* to set up dataset for analysis

## Import data from excel sheet
```{r import dataset}
#Import data
Dnet <- excel_to_network("data/DibamidsMouthBodyDemo.xlsx")
```

This is a list containing the graphs for 27 dibamid species.

## Construct distance matrix
The next step measures the distances and constructs the distance matrix.
It may take 10+ minutes. It saves the distance matrix to the "outputs" folder.

```{r measure distances, eval=FALSE, include=FALSE}
Ddist <- net_dist_mat(Dnet)

saveRDS(Ddist, file = "outputs/distance_matrix.rda")
```

For convenience(in case you're starting over from here), load in that distance matrix.

Next, we check the sums of the three distance measures, then combine the 3 distance measures into a single aggregate measure called comb.dist.We then chck for normality in the comb.dist distance matrix using a Shapiro-Wilks test.


```{r load distance matrix and analyze stats}
Ddist <- readRDS(file = "outputs/distance_matrix.rda")

sum(Ddist[,,1], na.rm = TRUE)
sum(Ddist[,,2], na.rm = TRUE)
sum(Ddist[,,3], na.rm = TRUE)

comb.dist <- (Ddist[,,3] + 
                Ddist[,,2] +
                Ddist[,,1])


sum(comb.dist, na.rm = TRUE)

median(comb.dist, na.rm = TRUE)
mean(comb.dist, na.rm = TRUE)

shapiro.test(comb.dist[lower.tri(comb.dist)])


```
Check whether the triangle inequality holds for comb.dist.

```{r check triangle inequality}
comb.dist[which(is.na(comb.dist))] <- 0
comb.dist <- comb.dist + t(comb.dist)

check.tri <- function(vec,distmatrix){
  
  distmatrix <- as.matrix(distmatrix)
  distmatrix[upper.tri(distmatrix)] <- t(distmatrix)[upper.tri(distmatrix)]
  a <- vec[1]
  b <- vec[2]
  c <- vec[3]
  distances <- c(distmatrix[a,b],
                 distmatrix[a,c],
                 distmatrix[b,c])
  ineqs <- c(sum(distances[1:2])>= distances[3],
             sum(distances[2:3])>= distances[1],
             sum(distances[c(1,3)]) >= distances[2])
  return(all(ineqs))
}

# test every
trieqs <- apply(combn(dim(comb.dist)[1],3), #all combinations of 2 dibamid species
                MARGIN = 2, check.tri, 
                distmatrix = comb.dist)

which(!trieqs)#nothing to see here! (the triangle inequality holds)

```

It does - we can't find any instances where it does not.

## Analyze the distance matrix and add other data

Since that's working, proceed to analysis with PCoA, & do k-means clustering.

```{r perform MDS on distance matrix}
PCoA <- cmdscale(comb.dist, eig=TRUE, k=2)
dimnames(PCoA$points)[[2]] <- c("PCoA 1","PCoA 2")
```

Combine the PCoA with traditional morphometrics and the following graph properties:
* density
* overall graph strength
* graph degree
* scale count
* composite scale count 
* number of fused edges
* number of fused scales

Run k-means clustering for k=2 & k=3 to understand whether geographic groups of Dibamus fall out differently.

```{r load in traditional morphometric data}
library('dplyr')
library('igraph')

D.data <- read.csv("data/D.data.csv")

D.data <- cbind(D.data, PCoA$points)

colnames(D.data)[8] <- "max.tl.proportion"
D.data$max.tl.proportion <- D.data$max.tl.proportion/100

D.data <- mutate(D.data,
                 density = vapply(Dnet, edge_density, numeric(1)),
                 strength = vapply(Dnet, FUN = function(x){sum(strength(x))}, 
                                   numeric(1)),
                 degree = vapply(Dnet, FUN = function(x){sum(igraph::degree(x))},
                                 numeric(1)),
                 scalecount = vapply(Dnet, FUN = function(x){length(V(x))},
                                     numeric(1))) %>%
  cbind(t(sapply(Dnet,
                 
                 function(g){
                   scalecount <- length(V(g))
                   el <- as_edgelist(g)
                   #what are the subgraphs of fused scales?
                   sg <- graph_from_edgelist(el[which(edge.attributes(g)$weight==2),]) 
                   
                   #how many subgraphs of fused scales are there, and how many 
                   comp <- components(sg)
                   
                   #return the original scalecount, minus all of the scales belonging to fused
                   # subgraphs, plus the number o ffused subgraph components
                   return(c("compScaleCount" = scalecount - sum(comp$csize) + comp$no,
                            "fusions" = sum(comp$csize),
                            "bigscales" = comp$no))
                 }
                 
  ))[,1:3])

D.data$k2 <- kmeans(scale(PCoA$points), centers = 2)$cluster %>% as.factor()
D.data$k3 <- kmeans(scale(PCoA$points), centers = 3)$cluster %>% as.factor()

colnames(D.data)[1] <- "label"

saveRDS(D.data, "outputs/Dibamusdf.rda")

```

# Analysis

## Visualize topospace
Plot the Principal coordinates space (Figure 3).
I use copwlot and ggpubr libraries here to quickly generate and arrange clean plots.

```{r plot PCoA space}
#Plots
library(cowplot)
library(ggpubr)
#colors
cl <- list("indo" = "#915062",
           "mx" = "#8EC260",
           "sea" = "#9BD2D4",
           "bg" = "#CED2D4",
           "cor" = "#187795")

hull <- D.data[,c("PCoA 1","PCoA 2","k2")] 
h1 <- chull(hull$`PCoA 1`[hull$k2 == 1],hull$`PCoA 2`[hull$k2 == 1])

h2 <- chull(hull$`PCoA 1`[hull$k2 == 2],hull$`PCoA 2`[hull$k2 == 2])

hulldf <- hull[c(c(which(hull$k2 == 1))[h1],c(which(hull$k2 == 2))[h2]),] 

hull <- hull[hull$edge,]

Fig.3 <- ggplot(data = D.data, aes(x = `PCoA 1`, y = `PCoA 2`)) +
  
  geom_point(aes(color = Locality, pch = Locality), size = 3) +
  scale_color_manual(values = c(cl$indo,cl$mx,cl$sea)) +
  #geom_label( aes(label = label,fill = Locality))+ #Turn this on to see species' positions
  xlab("PCoA 1")+
  ylab("PCoA 2")+
  theme_classic()+ 
  labs(tag = "A")+

  theme(legend.position = "bottom",
        panel.grid.major = element_line(colour = cl$bg),
        panel.grid.minor = element_line(colour = cl$bg, linetype = "dashed")) +
  geom_polygon(data = hulldf, alpha = 0.1, aes(fill = k2), show.legend = FALSE)
Fig.3
```

## Analyze relationships between topospace and characters

Can the peninsular/island and mainland groups be disrciminated from each other:

```{r discriminant function analysis}
install_github("ItziarI/WeDiBaDis")
library("WeDiBaDis")
discf <- WDBdisc(
  cbind(D.data$Locality,data.frame(comb.dist))[-1,-2] # remove Anelytropsis
  , datatype = "d", classcol = 1,
        method = "DB")
discf$conf
```
No, they cannot. 2 of the mainland species group with the Peninsular & Island species.

Plot correlations between principal coordinates and network and morphometric characters.

```{r plot correlations}

#Correlation Plots

scatterplot <- function(dataset,xvar,yvar, pos = "middle"){
  xpos <- min(dataset[,xvar])
  ypos <- min(dataset[,yvar])
  ggplot(data = dataset, aes(x = get(xvar), y = get(yvar))) +
    geom_smooth(method = "lm", color = cl$cor, fill = paste0(cl$bg,"05")) +
    geom_point(aes(color = Locality, pch = Genus), size = 3) +
    scale_color_manual(values = c(cl$indo,cl$mx,cl$sea)) +
    stat_cor(cor.coef.name = "R",
             label.x.npc = pos, 
             label.y.npc = "top",
             p.accuracy = 0.01,
             r.accuracy = 0.01,
             geom = "label")+
    xlab(xvar)+
    ylab(yvar)+
    theme_classic() +
    theme(legend.position = "none")
}

#plot size v TL and Component vs graph correlations.(Figure 4)

Fig.4 <- plot_grid(plot_grid(scatterplot(D.data,"scalecount","PCoA 1","left") + 
                      theme(axis.title.x=element_blank())+ labs(tag = "A"),
                    scatterplot(D.data,"fusions","PCoA 1","left") + 
                      theme(axis.title.x=element_blank())+ labs(tag = "B"),
                    scatterplot(D.data,"strength","PCoA 1","left") + 
                      theme(axis.title.x=element_blank())+ labs(tag = "C"),
                    scatterplot(D.data,"scalecount","PCoA 2","left") + 
                      xlab("scale count")+ labs(tag = "D"),
                    scatterplot(D.data,"fusions","PCoA 2","left") + labs(tag = "E"),
                    scatterplot(D.data,"strength","PCoA 2","left") + labs(tag = "F"),
                    ncol = 3),
          #scatterplot(D.data,"max.svl","max.tl.proportion","left") + labs(tag = "D"),
          #ncol = 1, nrow = 2, byrow = FALSE),
          get_legend(scatterplot(D.data,"max.tl.proportion","scalecount") + 
                       theme(legend.position = "bottom")), ncol = 1, rel_heights = c(1,.1))

#plot TL & SVL correlations (Figure 5)
Fig.5 <- plot_grid(
  plot_grid(scatterplot(D.data,"max.tl.proportion","scalecount")+
              ylab("scale count")+
              theme(axis.title.x=element_blank())+
              labs(tag = "A"),
            scatterplot(D.data,"max.tl.proportion","fusions")+
              theme(axis.title.x=element_blank())+
              labs(tag = "C"),
            scatterplot(D.data,"max.tl.proportion","PCoA 1")+
              theme(axis.title.x=element_blank())+
              ylab("PCoA 1")+
              labs(tag = "E"),
            scatterplot(D.data,"max.tl.proportion","PCoA 2")+
              ylab("PCoA 2")+
              xlab("maximum proportional tail length")+
              labs(tag = "G"),
            scatterplot(D.data,"max.svl","scalecount")+
              theme(axis.title.x=element_blank(), axis.title.y=element_blank())+
              labs(tag = "B"),
            scatterplot(D.data,"max.svl","fusions")+
              theme(axis.title.x=element_blank(), axis.title.y=element_blank())+
              labs(tag = "D"),
            scatterplot(D.data,"max.svl","PCoA 1")+
              theme(axis.title.x=element_blank(), axis.title.y=element_blank())+
              labs(tag = "F"),
            scatterplot(D.data,"max.svl","PCoA 2")+
              xlab("maximum SVL")+
              theme(axis.title.y=element_blank())+
              labs(tag = "H"),
            ncol = 2, nrow = 4, byrow = FALSE),
          get_legend(scatterplot(D.data,"max.tl.proportion","scalecount") + theme(legend.position = "bottom")), ncol = 1, rel_heights = c(1,.1))

Fig.4
Fig.5

```

Test models for Principal Coordinate Axes (Table 1). 

Rather than repeat a lot of code, I define a function, TL_models(), that runs all of the tests and then call it with different response variables.

I use the AICcmodavg library to produce AIC tables.

```{r PCo Models}
library(AICcmodavg)

TL_models <- function(df, col, PCoA = TRUE){
  models <- list(tl_str_loc = lm(eval(as.symbol(col)) ~ strength + Locality, data = df),
                 tl_scc_loc = lm(eval(as.symbol(col)) ~ compScaleCount + Locality, data = df),
                 tl_scl_loc = lm(eval(as.symbol(col)) ~ scalecount + Locality, data = df),
                 tl_fus_loc = lm(eval(as.symbol(col)) ~ fusions + Locality, data = df),
       tl_str = lm(eval(as.symbol(col)) ~ strength, data = df),
       tl_scc = lm(eval(as.symbol(col)) ~ compScaleCount, data = df),
       tl_scl = lm(eval(as.symbol(col)) ~ scalecount, data = df),
       tl_fus = lm(eval(as.symbol(col)) ~ fusions, data = df),
       tl_loc = lm(eval(as.symbol(col)) ~ Locality, data = df))
  
  nm <- c("strength + loc","compSc + loc","scalecount + loc","fusions + loc",
          "strength","compScaleCount","scalecount","fusions", "locality")
  if(PCoA){
    PCmodels <- list(tl_1_loc = lm(eval(as.symbol(col)) ~ `PCoA 1` + Locality, data = df),
                     tl_1 = lm(eval(as.symbol(col)) ~ `PCoA 1` , data = df),
                     tl_2_loc = lm(eval(as.symbol(col)) ~ `PCoA 2` + Locality, data = df),
                     tl_2 = lm(eval(as.symbol(col)) ~ `PCoA 2` , data = df))
    models <- append(models,PCmodels)
    nm <- append(nm,c("PCoA1 + Loc","PCoA1","PCoA2 + Loc","PCoA2"))
  }
  
  AIC = aictab(cand.set = models, 
                    modnames = nm)
  AIC$Rsq <- sapply(models, function(x){summary(x)$r.squared})
  AIC$adjRsq <- sapply(models, function(x){summary(x)$adj.r.squared})
  AIC$p <- sapply(models, function(x){summary(x)$p})
  return(list(models = models, AIC = AIC))
}

PCoA1_mod <- TL_models(D.data, 'PCoA 1',PCoA = F)
PCoA2_mod <- TL_models(D.data, 'PCoA 2',PCoA = F)
```

Test models for SVL and TL (Table 2), again using the TL_models() function.

```{r SV + TL models}
SVL_mod <- TL_models(D.data, 'max.svl')

TL_mod <- TL_models(D.data, 'max.tl.proportion')
```

Produce phylogenetic hypotheses.

Requires ape to handle trees, phytools to perfrom nj inference and plot the cophylo, and phangorn to perform parsimony inference.

```{r phylo}
library('ape')
library('phytools') 
library('phangorn')

Dchar <- morpho_matrix(Dnet)
# Dchar <- Dchar*2 - 1
# 
# Dchar[which(is.na(Dchar))] <- 0

D.phy <- nj(comb.dist)

contrast <- matrix(data = c(1,0,0,0,
                            0,1,0,0,
                            0,0,1,0,
                            0,0,0,1), ncol = 4, byrow = TRUE)
dimnames(contrast)<- list(c(0:3),c(0:3))
mm_pd <- phyDat(Dchar, type="USER", contrast=contrast)

mm_tree <- pratchet(mm_pd, start = random.addition(mm_pd), minit = 1000, maxit = 10000,
                    all = TRUE, trace = 0)
mm_tree <- acctran(mm_tree, mm_pd)
mm_cons <- consensus(mm_tree, p = 0.8)
mm_cons <- acctran(mm_cons, mm_pd)

cons <- CI(mm_cons,mm_pd)
ret <- RI(mm_cons,mm_pd)
hom <- 1-cons

par(mar = c(0,0,0,0))
Fig.6 <- plot(cophylo(mm_cons,D.phy), link.type = "curved",cex = 0.4)
Fig.6
```
````
