## Supplementary Tables, Figure S1 and Appendix for "A network-based comparative method to study reptile scalation and other homologous two-dimensional patterns"

| model | k | AICc | AICc Weight | Cumulative weight | Log Likelihood |
| --- | --- | --- | --- | --- | --- |
| PCoA 1 ~ strength | 3 | 111.73 | 0.89 | 0.89 | -52.34 |
| PCoA 1 ~ scale count | 3 | 117.36 | 0.05 | 0.94 | -55.16 |
| PCoA 1 ~ strength + locality | 5 | 117.48 | 0.05 | 0.99 | -52.31 |
| PCoA 1 ~ scale count + locality | 5 | 121.22 | 0.01 | 1.00 | -54.18 |
| PCoA 2 ~ fusions + locality | 5 | 117.17 | 1.00 | 1.00 | -49.16 |

*Table S1 – Most favored models for Principal Coordinate axes*

| model | k | AICc | AICc Weight | Cumulative weight | Log Likelihood |
| --- | --- | --- | --- | --- | --- |
| TL ~ PCoA 2 + locality | 5 | -106.74 | 0.28 | 0.28 | 59.80 |
| TL ~ composite scale count + locality | 5 | -106.66 | 0.27 | 0.54 | 59.76 |
| TL ~ fusions + locality | 5 | -105.84 | 0.18 | 0.72 | 59.35 |
| TL ~ strength + locality | 5 | -104.82 | 0.11 | 0.83 | 58.84 |
| TL ~ locality | 4 | -104.31 | 0.08 | 0.91 | 57.06 |
| SVL ~ PCoA 2 | 3 | 259.88 | 0.59 | 0.59 | -126.42 |
| SVL ~ composite scale count | 3 | 262.68 | 0.15 | 0.74 | -127.82 |
| SVL ~ PCoA 2 + locality | 5 | 264.11 | 0.09 | 0.83 | -125.62 |
| SVL ~ composite scale count + locality | 5 | 264.45 | 0.07 | 0.89 | -125.80 |
| SVL ~ fusions | 3 | 264.69 | 0.05 | 0.95 | -128.83 |

*Table S2 – Most favored models correlating maximum proportional tail length (TL) and snout vent length (SVL) to network summary statistics & principal coordinate values. Here, the locality of* Dibamus *species is used as a proxy for phylogenetic covariance.*

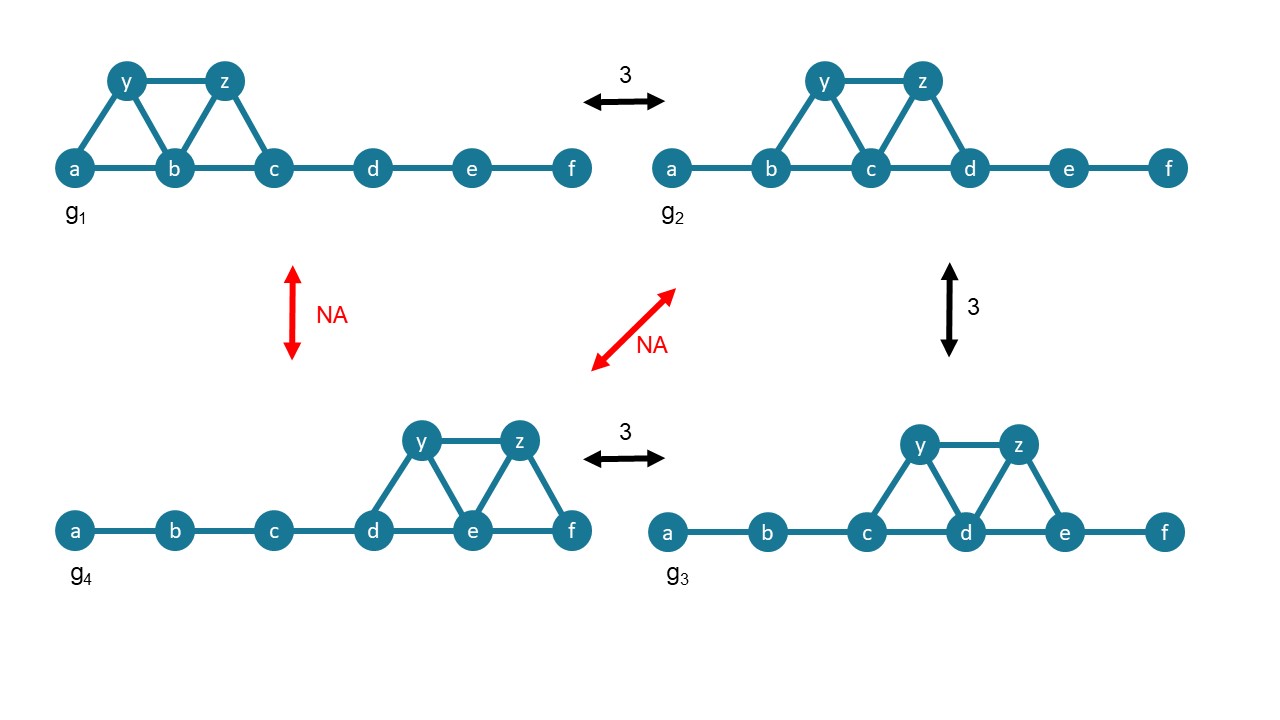

*Figure S1* – An example of topologies that cannot be compared using the algorithm described in this paper. While The graphs can be compared step-wise through this sequence of transformations, the algorithm described here cannot compare g_1_ to g_3_ or g_4_.

Appendix: References for Dibamid pholidosis networks

| Species | Reference |
| --- | --- |
| Anelytropsis papillosus | Greer 1985 |
| D. alfredi | Greer 1985 |
| D. bogadeki | Darevsky 1992 |
| D. booliati | Das & Yaakob 2003 |
| D. bourreti | Greer 1985 |
| D. celebensis | Greer 1985 |
| D. dalaiensis | Neang et al 2011 |
| D. deimontis | Kliukin et al. 2024 |
| D. deharvengi | Ineich 1999 |
| D. dezwani | Das & Lim 2005 |
| D. floweri | Quah et al. 2017 |
| D. greeri | Darevsky 1992 |
| D. ingeri | Das & Lim 2003 |
| D. kondaoensis | Honda et al. 2001 |
| D. leucurus | Greer 1985 |
| D. manadotuaensis | Koppetsch et al 2019 |
| D. montanus | Greer 1985 |
| D. nicobaricum | S.R. Chandramouli (pers. comm) |
| D. novaeguineae | Greer 1985 |
| D. seramensis | Greer 1985 |
| D. smithi | Greer 1985 |
| D. somsaki | Honda et al. 1997 |
| D. taylori | Greer 1985 |
| D. tebal | Das & Lim 2009 |
| D. tropcentr | Kliukin et al. 2023 |
| D. tiomanensis | Diaz et al. 2004 |
| D. vorisi | Das & Lim 2003 |
